# Crystal Structure of the Werner’s Syndrome Helicase

**DOI:** 10.1101/2020.05.04.075176

**Authors:** Joseph A. Newman, Angeline E. Gavard, Simone Lieb, Madhwesh C. Ravichandran, Katja Hauer, Patrick Werni, Leonhard Geist, Jark Böttcher, John. R. Engen, Klaus Rumpel, Matthias Samwer, Mark Petronczki, Opher Gileadi

## Abstract

Werner syndrome helicase (WRN) plays important roles in multiple pathways of DNA repair and the maintenance of genome integrity. Recently, loss of WRN was identified as a strong synthetic lethal interaction for microsatellite instable (MSI) cancers making WRN a promising drug target. Yet, structural information for the helicase domain is lacking, which prevents structure-based design of drug molecules. In this study, we show that ATP binding and hydrolysis in the helicase domain are required for genome integrity and viability of MSI cancer cells. We then determined the crystal structure of an ADP bound form of the WRN helicase core at 2.2 Å resolution. The structure features an atypical mode of nucleotide binding with extensive contacts formed by motif VI, which in turn defines the relative positioning of the two RecA like domains. The structure features a novel additional β-hairpin in the second RecA and an unusual helical hairpin in the Zn2+ binding domain, and modelling DNA substrates based on existing RecQ DNA complexes suggests roles for these features in the binding of alternative DNA structures. We have further analysed possible interfaces formed from the interactions between the HRDC domain and the helicase core by NMR. Together, this study will facilitate the structure-based design of inhibitors against WRN helicase.

## Introduction

Werner syndrome helicase (WRN) is one of the 5 human members of the RecQ family of DNA helicases that unwind DNA in a 3’ to 5’ direction and play important roles in multiple pathways of DNA repair and maintenance of genome integrity (1). Germline defects in three of these helicases lead to syndromes with hallmarks of premature ageing and cancer predisposition: Blooms syndrome (caused by mutations in Bloom’s syndrome helicase (BLM)), Rothmund-Thompson syndrome (caused by mutations in RECQL4) and Werner’s syndrome (WS). To date all WS mutations feature the introduction of premature stop codons or frame shift mutations that remove the nuclear localization signal at the C-terminus of WRN (2). Individuals affected by WS display many features associated with normal human ageing including premature greying and loss of hair, ocular cataracts, osteoporosis, atherosclerosis and an increased risk of development of various cancers. On a cellular level cells cultured from WS patients exhibit slow growth, chromosome aberrations, genome instability and an increased frequency of telomere shortening and loss, and are sensitive to various DNA damaging agents including radiation (3–5).

WRN is a 1432 amino acid, 162 kDa polypeptide that contains a central helicase core of 2 domains (D1 and D2) that share homology to *E. coli* RecA (residues 528-730 and 731-868 respectively), together with 3 additional helicase associated domains in the C-terminus: a Zinc binding subdomain (residues 869-994), a Winged Helix (WH) domain (residues 956-1064) and a Helicase and RNase D C-terminal (HRDC) domain (residues 1140-1239). WRN is unique among the RecQ family of helicases in containing a 3’-5’ exonuclease domain in the N-terminus (residues 38-236)(6). The nuclease domain has been characterized biochemically as being inactive on single stranded or blunt ended double stranded DNA, but capable of cleaving single nucleotides from the 3’ end of double stranded DNA containing 3’ recessed termini within the context of a variety of cellular DNA structures (7–9). Although WRN interacts with several proteins that participate in non-homologous end joining (NHEJ) (10,11), the role of WRN-nuclease activity in the context of DNA repair is poorly understood. WRN-deficient cells are unable to facilitate NHEJ-mediated double strand break repair (8). However, this deficiency can only be rescued by restoring both helicase and nuclease activities of WRN. Due to its interactions with various DNA repair proteins, WRN has also been implicated in several other cellular processes. During base excision repair, WRN interacts with DNA Polymerase beta wherein the helicase domain of WRN stimulates the strand displacement DNA synthesis activity of Polymerase beta. (12). In homologous recombination WRN interacts with RAD52 and increases its strand annealing activity; this interaction also modulates WRN helicase activity in a structure dependent manner (13). Rewrite Suggestion: Direct interactions of WRN with Replication protein A (RPA) (14–16), or shelterin complex components, TRF2 and POT1 have been shown to stimulate WRN helicase activity which is essential for DNA replication or Telomere maintenance respectively (17,18). Consistent with WRN playing a key role in telomere maintenance, cells from WS patients display telomerase dependent loss of telomeres from sister chromatids(3) and it has been speculated that the ability of WRN to unwind energetically stable non B-form DNA such as G-quadruplexes may explain this phenotype.

WRN was regarded as a ”Guardian of genome integrity” and a tumor suppressor. However, several recent studies have identified WRN as a potent and selective vulnerability of Microsatellite Instability-High (MSI-H) cancers (19–21). MSI-H cancers constitute a subset of colorectal, endometrial and gastric cancers that have deficiencies in the mismatch repair pathway and exhibit hyper-mutable state of microsatellite repeats. WRN was identified as the top dependency for MSI-H cells in two large genome wide gene inactivation studies using either CRISPR or RNA interference (19), and this dependency was linked to the helicase but not the nuclease function of WRN (19,20). These finding, together with the fact that WRN silencing in Microsatellite stable (MSS) cancer and normal cells is well tolerated, suggest that WRN is a promising novel drug target for the treatment of MSI cancers. To this end there have already been several high throughput screening efforts aimed at discovering potent and selective WRN inhibitors (22–24). And whilst these efforts appear to have produced compounds that induce DNA damage and apoptosis in cells(25), concerns about specificity, off target effects and cytotoxicity (20,26) indicate that significant improvements in potency and selectivity would be required to produce a pharmacologically useful compound.

In this study, we show that ATP-hydrolysis by WRN helicase is essential for maintaining cell viability and genomic integrity in MSI-H cells and determine the first crystal structure of the catalytic core of WRN helicase in complex with an ADP nucleotide at 2.2 Å resolution.

## Results and Discussion

### ATP-hydrolysis by WRN Helicase is essential for viability and genome integrity in MSI-H CRC cells

Previous studies(19–21,27) demonstrated that WRN depletion causes pervasive DNA damage and the loss of viability selectively in MSI-H but not MSS cell lines. Mutational analyses of WRN suggested that ATP-binding of the helicase domain but not enzymatic activity of the exonuclease domain is critical for the survival of MSI-H cells. It remained unknown if ATP-hydrolysis by WRN helicase and hence ATP turnover is required for the viability of MSI-H cells and which WRN enzymatic functions are essential for maintaining genomic integrity in MSI-H cells.

To address these questions,, we introduced the following mutations in WRN: the exonuclease-dead mutation E84A(9,28), the Walker A motif mutation K577M predicted to prevent ATP-binding, and the Walker B motif mutation E669A predicted to abolish ATP hydrolysis (28,29). We generated FLAG-tagged, siRNA resistant WRN (WRNr) transgenes that encoded wild-type or mutant variants of WRNr (Figure1A) and stably transduced them into the MSI-H colorectal cancer cell line HCT 116. Monoclonal HCT116 lines expressing the above WRNr variants were subsequently characterized by immunoblotting and immunofluorescence. We selected HCT 116 clones that showed expression and nuclear localization of the transgenic proteins (Figure S1A) and that exhibited transgene expression at a slightly higher level than the endogenous level of WRN protein (Figure 1B). Two different WRNr wild-type transgenic clones were selected based on their expression levels (WRNr wt high and low) to cover the range of protein expression levels displayed by the WRNr mutant transgenic clones. Depletion of the pan-essential mitotic kinase PLK1 by siRNA abrogated cell survival in all transgenic clones (Figure 1C). Upon depletion of endogenous WRN by siRNA, HCT 116 cells harbouring the empty vector lost cell viability, while the expression of wild-type WRNr at both low and high levels was able to restore viability (Figure 1C). This indicates on-target WRN knockdown and efficient transgenic complementation (Figure 1C). Consistent with previous reports (19–21,27), the WRNr E84A transgene also successfully rescued the WRN depletion viability phenotype (Figure 1C). Crucially, both the Walker A mutation K577M and the Walker B mutation E669A abrogated the WRNr transgene-mediated rescue of cell viability upon depletion of endogenous WRN (Figure 1C). This result strongly suggests that both ATP-binding and ATP-hydrolysis by the WRN helicase domain are essential for the survival of MSI-H cells.

**Figure 1.**
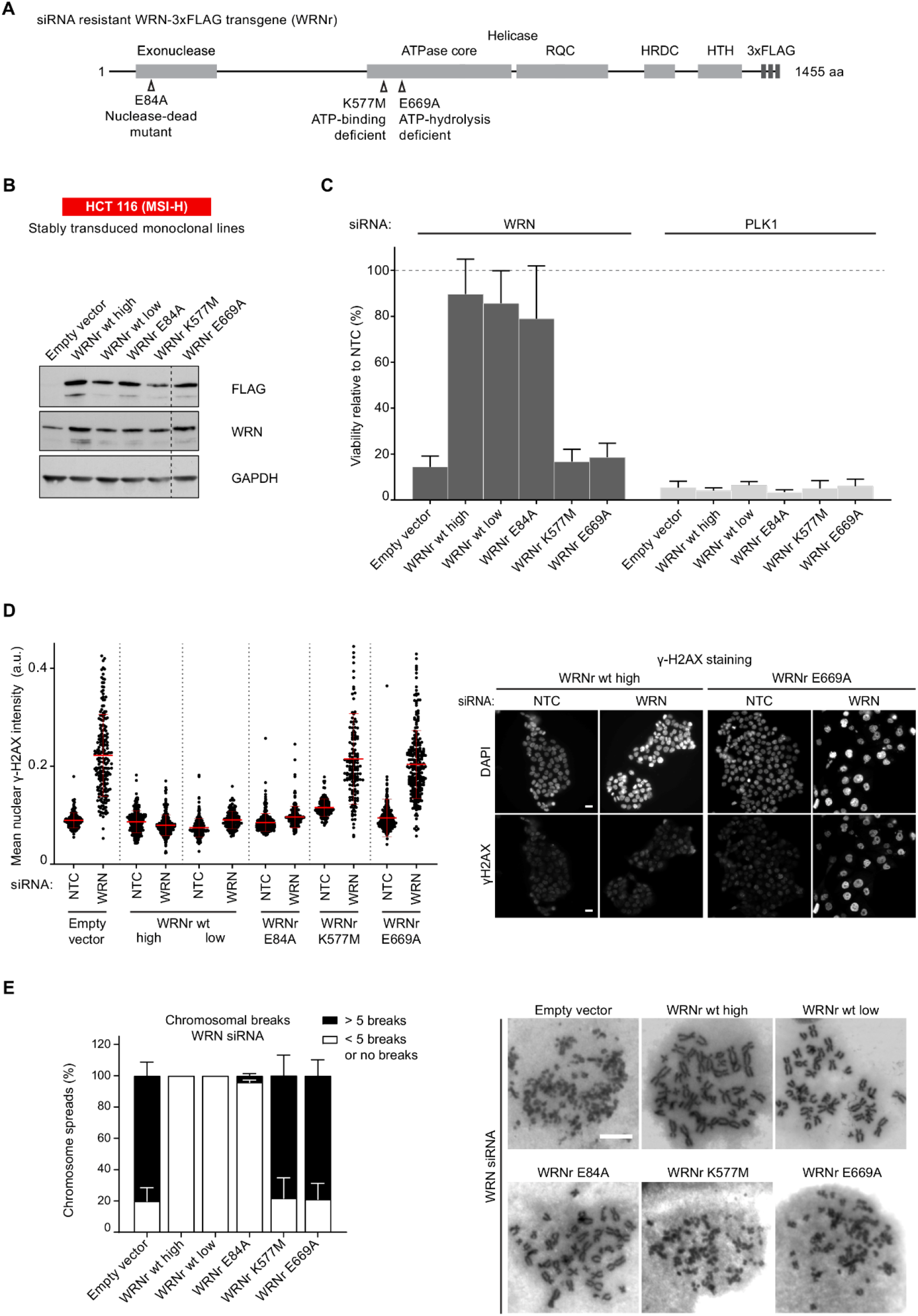
ATP hydrolysis by WRN is essential for essential for viability and genome integrity in MSI-H CRC cells. (**A**) Schematic representing the WRN domain structure. Location of nuclease-dead and ATPase-inactivating mutations (Walker A and B mutants) in siRNA-resistant WRN expression constructs containing a C-terminal 3xFLAG tag (WRNr) are indicated. (**B**) Monoclonal HCT 116 (MSI-H) cell clones were isolated after transduction with an empty vector control and WRNr wild-type or mutant transgenes. Immunoblotting of cell lysates with anti-FLAG and anti-WRN antibodies was used to determine the expression of the WRNr wild-type and mutant forms along with total WRN protein levels. Two WRNr wild-type clones (high and low) were selected to cover the expression range of WRNr mutant variants. (**C**) HCT 116 cells expressing WRNr transgenes were transfected with either NTC (non-targeting control) or WRN siRNAs. Viability measurements were performed 7 days after siRNA transfection and the data is represented relative to NTC siRNA. Data information: In (C), viability data are shown as mean ± SD of three biological repeat experiments. (**D**) Immunofluorescence analysis of γ-H2AX was performed 72 hr after siRNA transfection. The mean nuclear γH2AX intensity (a.u.-arbitrary units) was quantified after siRNA transfection. Data points shown (n ≥ 120 cells per condition) are derived from a single representative experiment that is consistent with a biological repeat experiment. Scale bar, 20 μM. (**E**) Mitotic chromosome spread analysis was performed 72 hr after siRNA transfection. At the 66 hr time-point, cells were treated with 6 hr of Nocodazole (1.5 μM) prior to spreading to enrich for mitotic stages. Each mitotic spread was categorized into less than 5 breaks or more than five breaks (n ≥ 28 mitotic spreads per condition). Data values and error bars presented here are the mean and the standard deviation respectively from biological repeats (n = 2). Scale bar, 10 μM

We next investigated the importance of specific enzymatic WRN functions for maintaining genome integrity in MSI-H cells. To this end, we performed immunofluorescence analyses of the DNA damage marker γ-H2AX and scrutinized chromosome breaks in mitotic spreads. siRNA depletion of WRN in cells harbouring an empty vector led to a strong increase in nuclear γ-H2AX and chromosome breaks. Expression of either the wild-type WRNr or the nuclease-dead WRNr E84A transgene prevented the increase in DNA damage markers. (Figure 1D and 1E; Supplemental Figure S1D and S1E). In striking contrast, siRNA depletion of endogenous WRN in cells expressing either the K577M or the E669A WRNr mutant transgenes displayed an increase in nuclear γ-H2AX levels and chromosome breaks comparable to WRN-depleted cells harbouring the empty vector (Figure 1D and 1E; Supplemental Figure S1D and S1E).

Our analyses of DNA damage markers and cell viability demonstrate that ATP-binding and ATP-hydrolysis by WRN helicase, but not WRN exonuclease activity, are essential for maintaining cell viability and genomic integrity in MSI-H cells. The genome stability defects observed across the enzymatic WRN mutants strongly correlate with the observed loss of viability phenotype. This suggests that pervasive DNA damage elicited by loss of WRN protein or loss of ATP turnover by WRN helicase is responsible for the attenuated viability of MSI-H cells. Further, our observations highlight the ATPase activity of WRN helicase as a therapeutic target to attack MSI-H cancers and provide pharmacodynamics markers of genome instability to track WRN ATPase inhibition in a cellular context.

### Crystal structure of the WRN ATPase core

Crystals of WRN were obtained using protein produced from overexpression in *E.coli* and a construct spanning residues 517-1093 with a C-terminal hexahistidine tag and TEV protease cleavage site. Crystallization was performed using sitting drop vapour diffusion and small shard like crystals appeared between one and two months. Crystals diffracted to 2.2 Å resolution and the structure was solved by molecular replacement using a domain based search strategy with the structure of Blooms syndrome helicase (PDBid 4CGZ) as a search model for the RecA domains and the WRN WH structure (PDBid 3AAF) as a search model for the WH domain. The crystallographic asymmetric unit contains a single molecule of WRN with no evidence for higher oligomer formation in the crystals. The model is well defined in the electron density with the exception of the first 12 residues in the N-terminus, a single loop spanning residues 950-953 and the final 21 residues in the C-terminus. The model has been restrained to standard bond lengths and angles with good geometry statistics (Table 1).

**Table 1.** Data collection and refinement statistics

|  | **WRN ADP** |
| --- | --- |
| Space group | P 2<sub>1</sub> 2<sub>1</sub> 2<sub>1</sub> |
| Cell dimensions | 54.6, 90.6, 138.2 |
| Angles $\alpha$, $\beta$, $\gamma$ (°) | 90, 90, 90 |
| Wavelength (Å) | 1.08 |
| Resolution (Å) | 76.0-2.20<br>(2.26-2.20) |
| *R*merge | 0.13 (2.3) |
| *R*<sub>p.i.m.</sub> | 0.06 (1.13) |
| *I*/σ*I* | 5.3 (1.0) |
| CC1/2 | 0.99 (0.50) |
| Completeness (%) | 98.8 (96.0) |
| Multiplicity | 5.5 (5.5) |
| No. of unique | 35356 (2943) |
| **Refinement Statistics** |  |
| Resolution | 76.0-2.20 |
| *R*work/*R*free (%) | 19.6/23.5 |
| **No. of atoms** |  |
| Protein | 4302 |
| Solvent | 195 |
| Ligand/Ion | 29 |
| **Average B factors (Å<sup>2</sup>)** |  |
| Protein | 71.8 |
| Solvent | 68.8 |
| Ligand/Ion | 76 |
| Wilson B | 48 |
| **RMSD** |  |
| Bond lengths (Å) | 0.002 |
| Bond angles (°) | 0.534 |
| **Ramachandran plot** |  |
| Favoured (%) | 97.3 |
| Allowed (%) | 2.7 |

The overall structure of WRN consists of two RecA like helicase lobes D1 and D2, each featuring a central 6 stranded parallel β-sheet flanked on each side by helices and loops (Figure 2A). The Zn^2+^ binding subdomain features a single Zn ion tetrahedrally coordinated by four cystine residues (C908, C935, C936 and C939) and is closely associated with the D2 domain. The WH domain extends away from the Zn^2+^ and D2 domains and features a modified version of the canonical WH fold with a longer wing 2.

**Figure 2.**
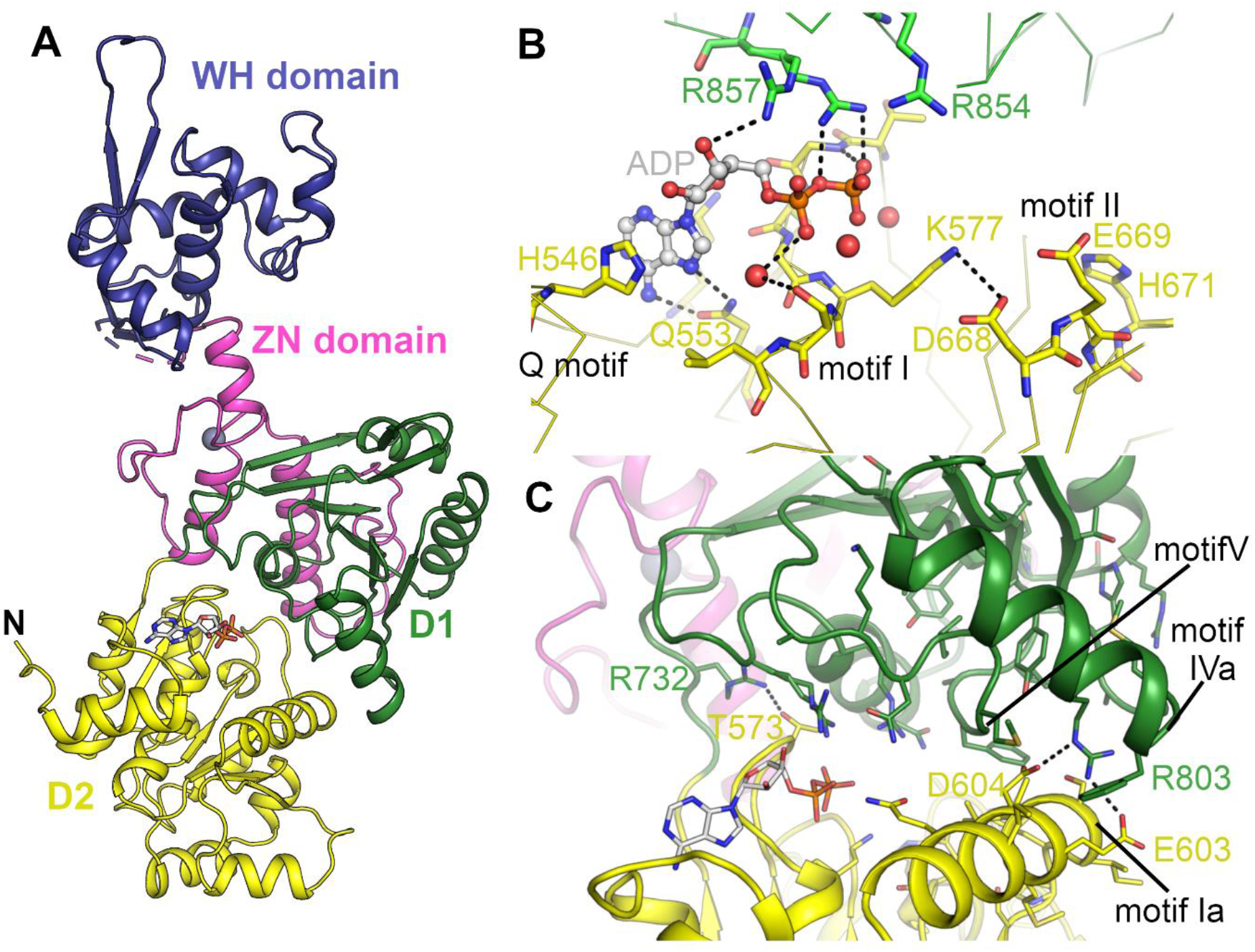
Structure of WRN helicase and the nucleotide binding site. **A** Overall structure of WRN helicase with domains coloured individually. **B** Close up view of the WRN nucleotide binding site with conserved helicase motifs and key residues labelled. **C** Close up view of the contact formed and interface between the D1 and D2 domains with key residues and motifs labelled.

### Structure of the nucleotide binding site

The nucleotide binding site is positioned in the cleft between D1 and D2 with the majority of contacts to the nucleotide coming from D1. The protein was crystallized in the presence of ADP, Mg^2+^, aluminium chloride and sodium fluoride, intended to produce the ATP analogue ADP-Aluminium fluoride, although examination of the electron density reveals only density for ADP; similarly, no convincing electron density could be observed for the magnesium ion. The adenine moiety on the ADP is flanked on either side by H546 and K550 and forms polar contacts with Q553, part of the conserved Q motif common to all RecQ family members (Figure 2B). The ribose makes a single contact to R857 and the phosphates are positioned directly above the N-terminal end of helix α3 within the motif I or Walker A motif. In contrast to what has been observed for other RecQ family member structures, the catalytically essential K577 does not form direct hydrogen bonds to the β-phosphate, instead forming polar contacts with motif II (Walker B motif). This is also true of other residues within motif I which generally make less direct and more water mediated contacts with the phosphates than what has been observed in previous RecQ family structures (Figure S2). On the other hand, the contacts made by residues belonging to D2 are more extensive than that observed in other RecQ structures. R857, one of two highly conserved arginines from motif VI, the so called “arginine finger” shows a dual conformation in which it makes contacts with both ribose and phosphates. R854, the first conserved arginine from this motif, is in a position to interact with the expected location of the γ-phosphate in an ATP molecule (Figure 2B). This means of contacting the nucleotide, with potentially both conserved arginine residues, has not been seen in other RecQ family structures to date. A mutational analysis in Blooms syndrome helicase (BLM) indicated that both residues are important for helicase activity with the equivalent of R857 identified as the transition state stabilizing arginine finger(30).

### Comparisons with other RecQ family members

The extensive contacts formed between R857 and the ADP nucleotide play a role in defining the relative positioning of the two RecA-like domains which we and others have found to be variable in RecQ family structures. We have previously performed a systematic analysis of domain positioning in existing RecQ family structures by measuring vector pairs between invariant points on each RecQ domain(31). Using the same analysis on the WRN structure reveals that the positioning of the two RecA domains in WRN is unique (Figure S3) with a significantly more compact arrangements of the two domains, and close contacts formed between motif Ia in D1 and motifs IVa and V in D2, (Figure 2C). On an individual domain basis the WRN structure is surprisingly most similar to RecQ from *Deinococcus radiodurans*(32) (around 1.6 Å RMSD for both D1 and D2), although the similarity to other human RecQ family members is only slightly lower (generally around 1.8 Å RMSD) (Figure 2A). One unique feature of the WRN is an additional β-hairpin inserted between the first helix and second strand of the D2 domain (Figure 3A), the hairpin is highly reminiscent of the strand separating hairpins found in various helicases, and features a compact type II’ β-turn with a serine (S758) instead of the usual glycine residue at the +1 position. Similar hairpin features have been observed as additions to the second RecA domain in other helicases such as the superfamily I PcrA and superfamily II Hel308 (33,34), although in the case of WRN the inserted hairpin is on the same face but opposite ends of the D2 domain. The amino acid sequence at this region is not well conserved in WRN homologues although this is also the case for the canonical strand separating hairpin (aa 1028-1043) in the WRN WH domain and in hairpins found in helicases from other organisms.

**Figure 3.**
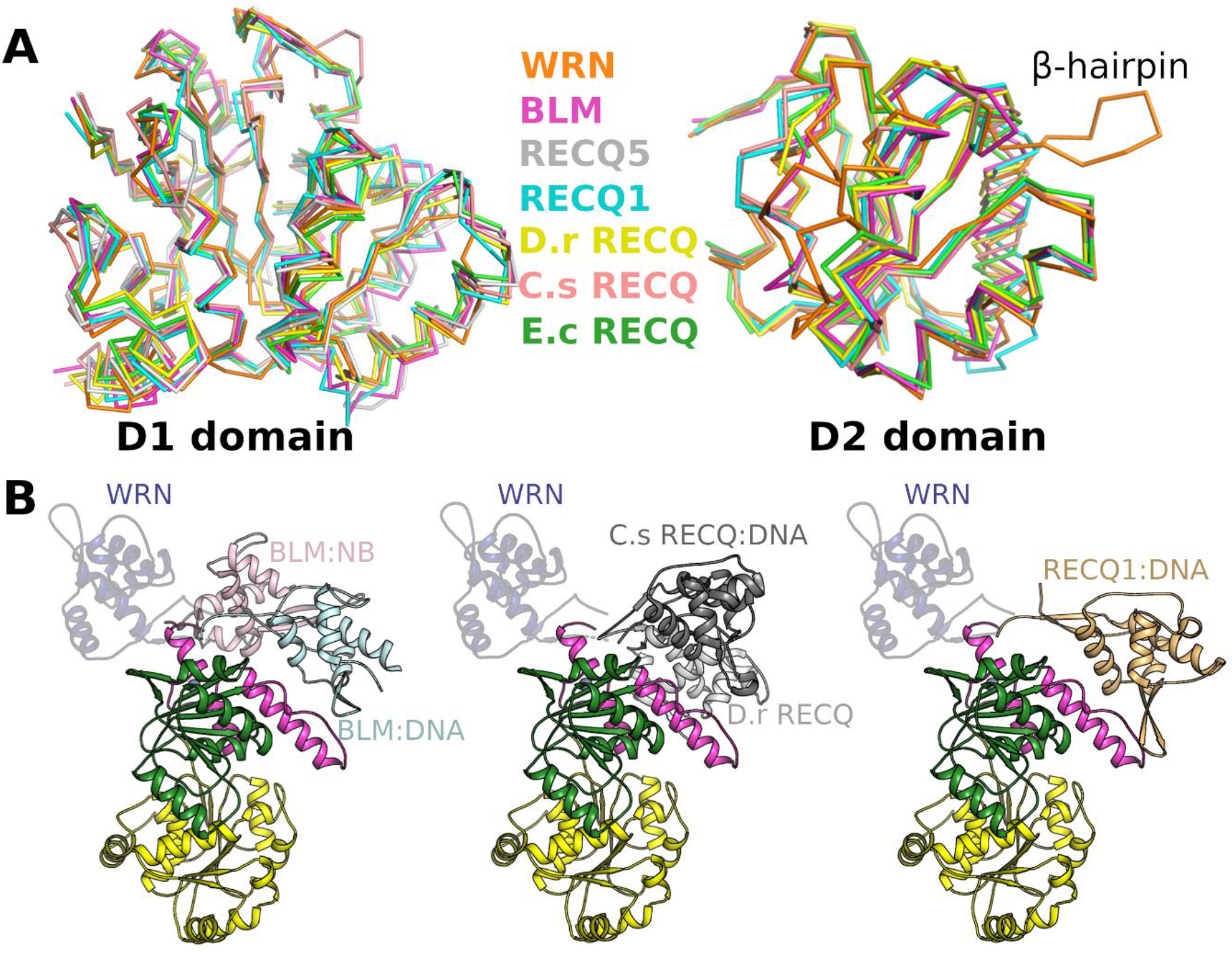
Comparison of WRN with other members of the RecQ family. **A** Comparison of current RecQ helicase structures superposed on the basis of the D1 domain (left) and D2 domain (right). **B** Comparison of relative positioning of the WH domain with respect to the helicase core in various RecQ family structures, alignments were performed on the basis of the D2 domain with BLM nanobody complex vs BLM DNA shown on the left, Bacterial RecQ APO vs Bacterial RecQ DNA in the centre and RecQ1 shown on the right. The WRN WH domain is shown throughout in semi-transparent blue.

Another notable difference in the WRN structure is the position adopted by the WH domain relative to the D2 domain. The positioning of this domain has been found to vary in other RecQ helicase structures, with the positioning in the absence of DNA being quite varied, whilst the DNA complexes are more consistent, with the WH domain packed closely against the helical hairpin of the Zn domain(35,36). In the WRN structure the WH domain is positioned in an opposite orientation and displaced by around 30 Å from the typical WH positioning in RecQ DNA complexes (Figure 3B). The presence of numerous crystal contacts and an intra-molecular disulphide bond (C946 to C1070) indicate that this positioning is not expected to be representative of the DNA bound conformation, although the flexible attachment of this domain may be a feature of the WRN helicase mechanism and alternative conformations may be required for activity on unusual DNA substrates.

### Potential interactions of the WRN HRDC domain with the helicase core

The function of the HRDC domain in various RecQ helicases is currently an active area of research. Early studies on *E. coli* and *S. cerevisiae* RecQ proteins indicated a role as an accessory DNA binding domain(37,38), with the isolated HRDC domains having an electropositive surface and displaying binding affinity for ssDNA in the low to mid micro molar range(38,39). On the other hand structural studies on isolated HRDC domains from BLM and WRN did not find an electropositive surface and no DNA binding activity could be found for WRN HRDC, although BLM HRDC was reported to have weak ssDNA binding affinity (^~^100 μM) in one study, whilst another failed to detect any binding at all(40–42). Further clues as to the role of the HRDC domain came from structural investigations of human BLM which showed that in the presence or absence of DNA the HRDC domain packs tightly against the helicase core and forms interactions with both D1 and D2 domains in a nucleotide dependant manner (35,43). Subsequent studies with *E. coli* RecQ showed the HRDC domain suppresses the rate of ATP hydrolysis and DNA unwinding independently of its ability to bind DNA(44). From this it has been suggested that interactions between the helicase core and HRDC domains is a conserved feature of RecQ helicases, although the effect on the helicase activity may vary according to the roles of the different enzymes.

To investigate this possibility for WRN, we have constructed a model of the possible interface between the HRDC domain and the helicase core using the structure of the WRN HRDC domain determined in isolation together with the WRN helicase core and the relative domain positioning found in the BLM helicase structures (35,41). In this model there is generally good shape complementarity between the WRN HRDC domain and its expected interface (Figure 2B), with some minor clashes formed by hydrophobic residues at the C-terminal end of the first helix of the HRDC domain that can be largely relieved by adopting alternative rotamers for the affected residues. The putative interface in the WRN structure is slightly smaller and less polar than in the BLM structure,with significantly fewer salt bridges (1 versus 8) and hydrogen bonds formed (7 versus 17). Nevertheless the WRN HRDC can be seen to make potentially favourable pairs of interactions to D1 (primarily hydrophobic in nature) and D2 (more polar), with the nucleotide being in close proximity to a number of polar residues in the interface K1182 and T1180 (Figure 4A). Other unique features of the WRN HRDC domain such as the extended N-terminal helix and C-terminal loop motif (41) are found on the opposite face to the expected interface suggesting an interaction is possible. We have tested this possibility in solution by NMR using a ^15^N labelled HRDC domain expressed separately. As can be seen in Figure 4B, no chemical shift perturbations can be observed after the addition of WRN Helicase, indicating that there is no interaction between the unconnected HRDC- and helicase domains of WRN under the experimental conditions. This indicates that potential interactions of the HRDC with the helicase core are significantly less extensive than that found for BLM. Additionally, the linker region between the end of the WH domain and the start of the HRDC is significantly longer in WRN (around 70 residues) than in BLM (around 10 residues). This additional flexibility in combination with a less extensive interaction network may hamper a strong interaction between these domains, whilst enabling the WRN HRDC module to play additional roles.

**Figure 4.**
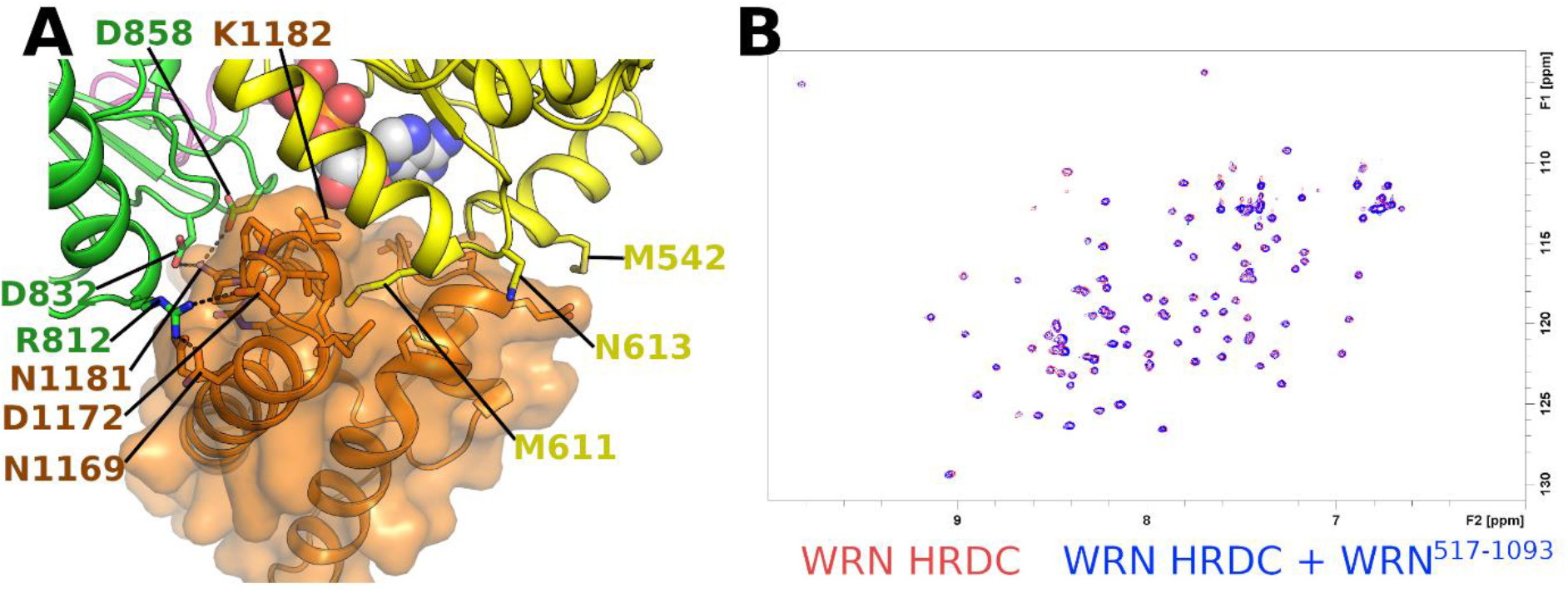
Examination of possible contacts between the WRN HRDC domain and helicase core. **A** structural model of the possible WRN HRDC domain – helicase core interaction interface created by positioning the isolated WRN HRDC structure into its expected position based on the BLM helicase structure. **B** Superposition of the 2D 15N SoFast HMQC spectra of 10 uM 15N-labeled WRN HRDC in absence (red) or presence (blue) of 20 μM unlabelled WRN Helicase (residues 517-1093)

### A model for WRN DNA binding

We have also used the existing available structural information to construct a model for WRN helicase bound to a simple DNA substrate (Figure 5A). The model is constructed by positioning the WRN WH domain onto the position adopted by the *Chronobacter sakazakii* RecQ-DNA complex (36). This model was chosen as the WH domain positioning in either the BLM or RecQ1 DNA bound structures whilst being broadly similar give a significant number of steric clashes due to the unusual conformation adopted by the WRN helical hairpin. The double stranded DNA from the WRN WH DNA complex structure was extended to include a 4 nucleotide 3’ overhang with the positioning of the 3^rd^ and 4^th^ nucleotides of the overhang in close contacts with residues from conserved helicase motifs IV and V respectively, a feature common to all RecQ DNA structures determined to date (Figure S4). The model is completed with an additional two unpaired bases connecting these two DNA elements with the conformation of these nucleotides being similar to that observed in the RecQ1 DNA structure (45) although there is significantly more uncertainty over this region as it is quite variable across the various DNA bound RecQ structures determined to date.

**Figure 5.**
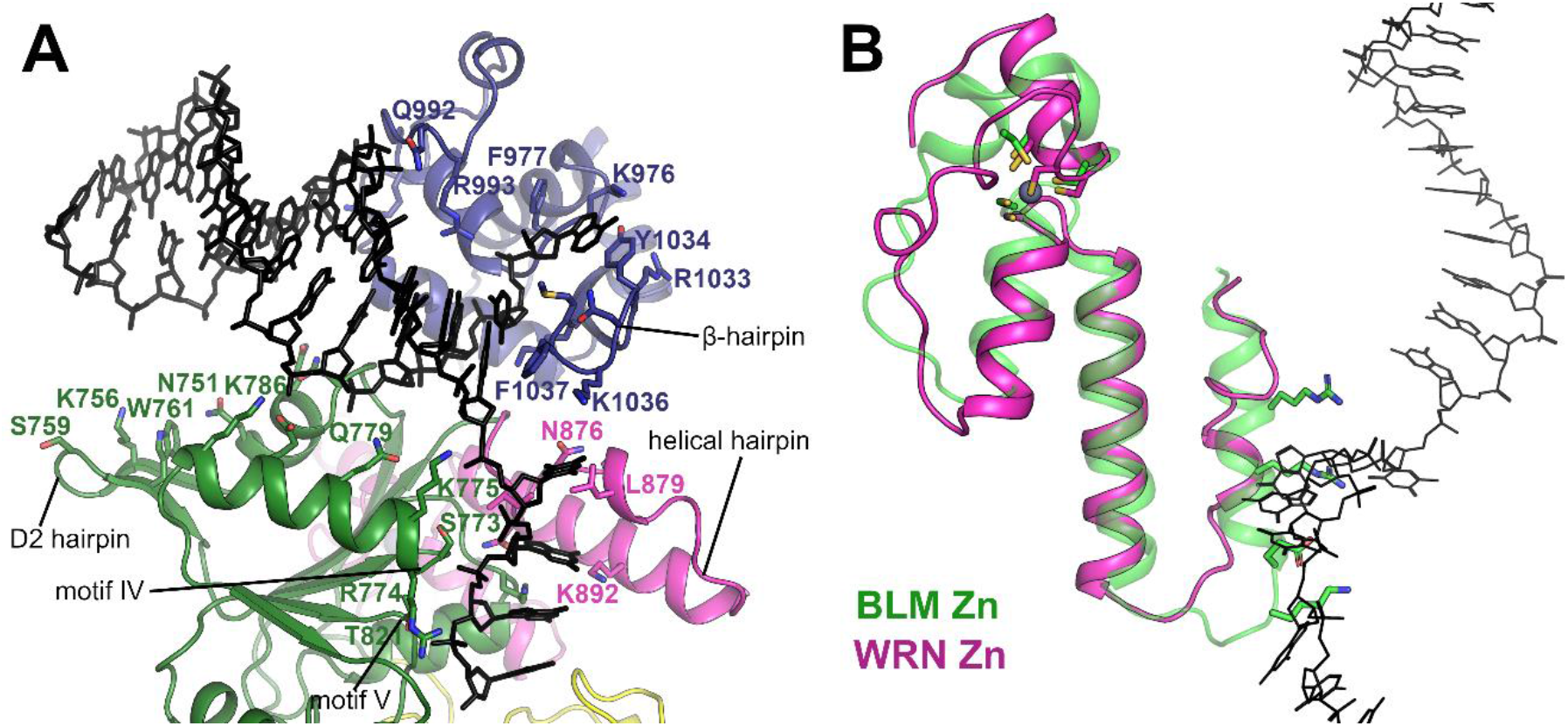
A model of WRN bound to DNA containing a 3’ overhang. **A** Overview of the WRN DNA model with predicted DNA contacting residues and motifs labelled. **B** Comparison of the Zinc binding domain in BLM (green) and WRN (pink) helicases and its contacts to the 3’ DNA overhang (shown in black stick format), the WRN Zn binding domain features an extended linker helix, alternate positioning of Zn coordinating residues, and coil conformation of the N-terminal arm of the helical hairpin. Sidechain residues from the helical hairpin that form contacts to DNA in the BLM structure are shown in stick format for reference.

In the model the double stranded region of the DNA sits in a cleft between the D2 and WH domains and makes extensive interactions with the WH domain in addition to potential favourable interactions formed by polar residues on the 2^nd^ helix in D2 such as K775, Q779 and K786. The blunt end of the double stranded DNA is also close to the WRN specific hairpin insertion on the D2 domain and a number of polar residues from this and the preceding α-helix point towards the DNA duplex and are suggestive of a possible role in the protein DNA interface, although in the current model DNA is slightly too distant for formation of direct contacts. The single stranded 3’ overhang passes close to the helical hairpin region of the Zn domain which in WRN is distinctly different from that found in all other RecQ structures, with the N-terminal helix being predominantly coil rather than helical (Figure 5B). This helix forms significant contacts to the single stranded DNA in the other RecQ structures, generally in the form of hydrophobic contacts from bulky side chain residues contacting the nucleobases, whilst in the WRN DNA model the equivalent residues are more distant with significantly more room for the DNA to pass unhindered, and additional pockets on the surface created by the helix to coil transition (Figure 5B). One clue as to the possible function of such a structure came from a recent structural study on *C. sakazaki* RecQ, in complex with an unwound G-quadruplex DNA, which found a guanine specific pocket that accommodated a flipped out Guanine nucleobase with residues in this pocket being identified as essential for G4 unwinding(46). The pocket identified for bacterial RecQ is not conserved in WRN, although it may be possible that the pockets formed by the helix coil transition of the helical hairpin, which are in a similar position but on the opposite side of the DNA tract, may play the same role.

### Hydrogen Deuterium Exchange (HDX) measurements of WRN in solution

We have probed the WRN interaction with both nucleotide and DNA in solution by performing HDX MS measurements of WRN in the presence and absence of both ssDNA and the non-hydrolysable ATP analogue AMPPNP using a shorter construct (including residues 531 to 950) lacking the WRN WH domain, which gave excellent peptide coverage of 98 %. As can be seen in Figure 6A, a significant protection of residues close to the Q motif, and motifs I and III can be seen in the presence of nucleotide. In particular the peptide spanning residues 571-580 containing motif I shows a reduced deuteration of between 2-3 Daltons, which is consistent with the typical mode of nucleotide interaction with motif I, with 3 consecutive direct hydrogen bonds donated by backbone amides, as found in other RecQ nucleotide structures (Figure S1). With longer incubation periods (4 hours) additional protection can be seen for residues from motif VI in the D2 domain (Figure 6A), which is consistent with the extensive interactions between that region and the nucleotide in our crystal structure. Addition of single stranded DNA alone does not cause any differences in exchange at short labelling times, however after 4 hours of labelling, significant protection can be seen for a peptide containing part of the D2 hairpin and the entire helicase motif IV (Figure 6B), although the sequence coverage for this measurement was significantly lower (73%). Both the D2 hairpin and motif IV are predicted in our WRN DNA complex model to have the potential to interact with DNA, therefore indicating that the HDX data support our modelling studies. These HDX results further suggest that the compact arrangement of D1 and D2 domains found in our crystal structure, and the extensive contacts formed between nucleotide and D2, may be a feature of the WRN protein in solution. The unusual mode of nucleotide interaction with motif I seen in the crystal structure does not appear to be predominant in solution, as revealed by HDX, and thus may be associated with a particular WRN conformational state rather than being a general feature of the WRN protein.

**Figure 6.**
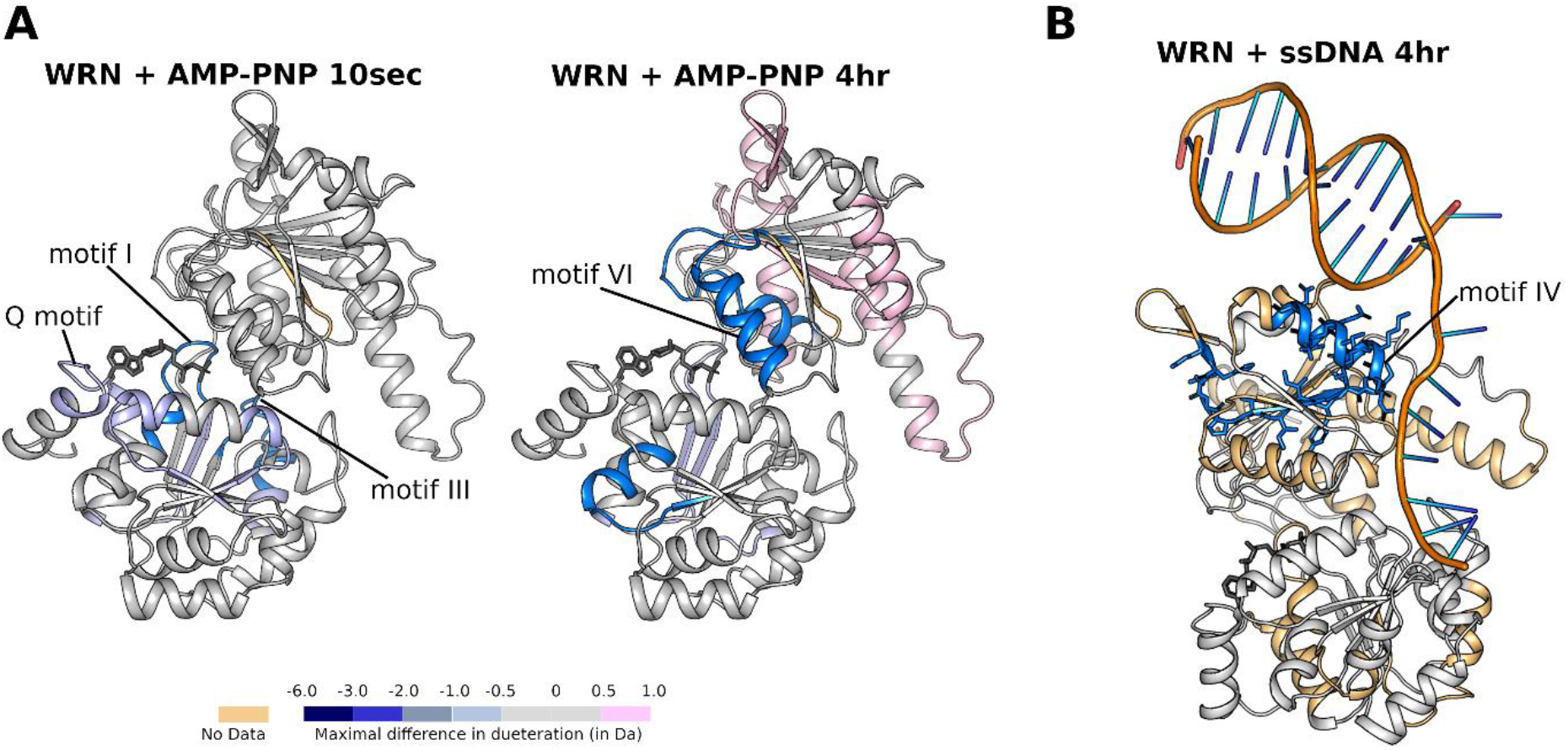
Hydrogen deuterium exchange MS measurements of WRN in solution. **A** Comparative HDX (D_AMP-PNP_ – D_unbound_) of WRN in complex with the ATP analogue AMP-PNP mapped onto the WRN structure. Protection can be seen for nucleotide binding features in D1 (left), whilst protection for nucleotide contacting residues from D2 are observed over longer time scales (right). **B** Comparative HDX of WRN (D_ssDNA_ – D_unbound_) in complex with single stranded DNA, mapped on to the WRN DNA binding model. Protection can be seen for residues in the D2 hairpin and motif IV.

## Summary

We have determined the crystal structure of WRN helicase, a highly anticipated structure due to the recently discovered importance of WRN as selective dependency of and therapeutic target in MSI cancer cells. The structure shows an unusual mode of nucleotide binding where interactions formed by D2 are more extensive than that found in previous RecQ family structures whilst those formed by the walker A motif in D1 are less so. This interaction with the nucleotide in turn defines the relative domain positioning of the D1 and D2 domains which form a compact arrangement distinct from that seen in other RecQ structures. The WRN WH domain adopts a conformation that appears to not be compatible with DNA binding, although this has been observed for other RecQ structures in the absence of DNA, and we have used this similarity to construct a model for WRN DNA binding which suggests possible roles for a WRN specific insertion in the D2 domain and an unusual helical hairpin in defining the DNA protein interface.

## Experimental procedures

### Cell culture and lentiviral transduction

The human colon cancer cell line HCT 116 was grown in McCoy’s 5A medium with glutamax (GIBCO, 36600–021) to which 10% fetal calf serum (FCS) was added. Wild-type and mutant codon-optimized, siRNA resistant WRN transgenes containing a C-terminal 3xFLAG tag (designated WRNr) were synthesized and inserted into the lentiviral pLVX-IRES-puro plasmid vector (ClonTech) at GenScript. Lentivirus particles were generated using the Lenti-X Single Shot system (ClonTech, Mountain View, CA, US) in 293T-Lenti-X cells. HCT 116 cell pools stably transduced with the WRNr transgene carrying lentivirus particles were selected with 2 μg/ml of Puromycin (SIGMA, P9620) added to the normal growth medium. These cell lines were generated using lentiviral transduction. Single cell clones were obtained by limiting dilution. All cell lines used in this study tested negatively for mycoplasma contamination and were further authenticated by STR fingerprinting.

### Immunoblotting

Cells were lysed in extraction buffer (50 mM Tris-HCl pH 8.0, 1% Nonidet P-40, and 150 mM NaCl) to which Complete protease inhibitor mix (Roche, Switzerland) and Phosphatase inhibitor cocktails (SIGMA, P5726 and P0044) were added.

### Antibodies

The antibodies used in this study are as follows: WRN (8H3) mouse mAb (Cell Signaling, 4666, 1/1000 dilution), mouse anti-FLAG (SIGMA, F1804, 1/1000 [immunoblotting] or 1/500 [immunofluorescence] dilution), rabbit anti phospho-Histone H2A.X (Ser139) (Cell Signaling, 2577, 1/800 dilution), mouse anti-GAPDH (Abcam, ab8245, 1/30000 dilution), mouse Alexa Fluor 488 (Molecular Probes, Eugene, OR, US, 1/1000 dilution) and secondary rabbit (Dako, P0448, 1/1000 dilution), mouse anti-IgG-HRP (Dako, P0161, 1/1000 dilution)

### siRNA transfection and cell viability

For siRNA knock-down experiments, cells were transfected using Lipofectamine RNAiMAX reagent according to the manufacturer’s instructions (Invitrogen, Waltham, MA, US) supplemented with the following siRNA duplexes: WRN and PLK1 targeting ON-TARGETplus siRNA duplex (J-010378–05, L-003290-00, Dharmacon, Lafayette, CO, US); ON-TARGETplus Non-targeting Control (NTC) Pool (D-001810-10, Dharmacon, Lafayette, CO, US). The final concentration of the siRNA was 20 nM in immunoblotting, immunofluorescence and chromosome spread experiments. Cell viability experiments were carried out with 10 nM siRNA concentration in 96-well plates with a total volume of 100 μl per well and a starting number of 1000 HCT 116 cells per well. Cellular viability was measured seven days after transfection using CellTiter-Glo reagent (Promega, Madison, WI, US). 100 μl of the 1:2 diluted CellTiter-Glo solution was directly added to the growth medium, mixed briefly and incubated for 10 min prior to the measurement of the luminescence signal.

### Immunofluorescence

Cells were transfected as mentioned above and grown for 72 hrs. Subsequently the cells were fixed for 15 min using 4% paraformaldehyde, permeabilized for 10 min with 0.2% Triton X-100 in PBS and blocked for 45 min with 3% BSA in PBST (PBS containing 0.01% Triton X-100). Cells were incubated sequentially with primary antibodies that detect either FLAG or phospho-Histone H2A.X (Ser139) and secondary antibodies (Alexa 488, Molecular Probes, Eugene, OR, US). Coverslips were mounted and cells were counterstained on the glass slides using ProLong Gold with DAPI (4’, 6-diamidino-2-phenylindole) (Molecular Probes, Eugene, OR, US). Images were collected using an Axio Plan2/AxioCam microscope and image processing was performed with MrC5/Axiovision software (Zeiss, Germany). Quantification of γ-H2AX foci was carried out using segmentation in the Halo software (https://www.indicalab.com/halo/) that identified DAPI-stained nuclei. Subsequently, the corresponding γ-H2AX mean intensities of the identified nuclei were determined.

### Chromosome spreads

66 hr after siRNA transfection, cells were treated with 1.5 μM Nocodazole for 6 hr. Cells were swollen in hypotonic buffer for 5 min at room temperature in a solution composed of 40% medium/60% tap water. Fixation was performed three times with freshly made Carnoy’s solution (75% methanol, 25% acetic acid). To acquire chromosome spreads, cells in the fixative solution were dropped onto glass slides and air dried. Slides were later stained with 5% Giemsa (Merck) for 4 min, washed briefly with tap water and air-dried. The analysis was performed from two independent slides for each condition and a blind quantification of the chromosome breaks was carried out. Images were acquired using an Axio Plan2/AxioCam microscope and image processing was performed with MrC5/Axiovision software (Zeiss, Germany).

### Cloning, Overexpression and Purification of WRN helicase domain

WRN constructs corresponding to residues 517-1093 were cloned in the vector pNIC28-Bsa4 using ligation independent cloning and transformed into E. coli LOBSTR cells for overexpression(47). Cells were grown at 37°C in TB medium supplemented with 50 ug/ml kanamycin until an optical density of 2-3 and induced by the addition of 0.1 mM IPTG and incubated overnight at 18°C. Cells were harvested by centrifugation. For purification, cell pellets were thawed and resuspended in buffer A (50 mM HEPES pH 7.5, 500 mM NaCl, 5% glycerol, 30 mM imidazole, 0.5 mM Tris (2-carboxyethyl) phosphine (TCEP)), with the addition of 1x protease inhibitor set VII (Merck, Darmstadt, Germany). Cells were lysed by sonication and cell debris pelleted by centrifugation. Lysates were loaded on to a Ni-sepharose IMAC gravity flow column (GE healthcare), washed with 2 column volumes of wash buffer (buffer A supplemented with 45 mM imidazole), and eluted with 300 mM imidazole in buffer A. The purification tag was cleaved with the addition of 1:20 mass ratio of His-tagged TEV protease during overnight dialysis into buffer A. TEV was removed by IMAC column rebinding and flow through and wash fractions were combined, concentrated using a 50,000 mwco centrifugal concentrator and loaded on to size exclusion chromatography using a HiLoad 16/60 Superdex s75 column in buffer A. Fractions containing WRN were pooled, and diluted to 25mM Hepes, 250 mM NaCl, 2.5 % glycerol, 0.25 mM TCEP and loaded onto a 1ml HiTrap Heparin HP column, equilibrated in the same buffer. Proteins were eluted with a 40 ml linear gradient to 50 mM Hepes, 1 M NaCl. Protein concentrations were determined by measurement at 280nm (Nanodrop) using the calculated molecular mass and extinction coefficients.

### Crystallization and Structure Determination

For crystallization WRN was concentrated to 12 mg/ml using a 50,000 mwco centrifugal concentrator diluted 2 fold in water and co crystallized with 5mM AlCl3, 60 mM NaF, 5 mM ADP, 5 mM MgCl at a final protein concentration of 5.5 mg/ml. WRN crystals appeared between 1 and 2 months in conditions containing 1M Na Acetate, 0.1 M Cacodylate pH 6.5. Crystals were cryo-protected by transferring to a solution of mother liquor supplemented with 20 % Glyecrol and flash-cooled in liquid nitrogen. Data were collected at Diamond Light Source beamline I03, and data were processed with the programs DIALS(48). The structure was solved by molecular replacement using the program PHASER(49) with the RECQL1 structure as a starting model(45). Model building and real space refinement were performed in COOT(50) and the structures refined using PHENIX REFINE(51). A summary of the data collection and refinement statistics is shown in Table I.

### Expression and Purification of ^15^N-labeled WRN HRDC Domain

The construct for expression of the WRN HRDC domain (residues 1142–1242), as described previously (41), was obtained by gene synthesis (GeneArt, Thermo-Fisher) in a donor vector (pDONR-221) and transferred by recombinant cloning into the glutathione S-transferase (GST) fusion vector pDEST15 (Invitrogen). The plasmid was used to transform Escherichia coli, strain BL21(DE3). An overnight culture in LB-media supplemented with 100μg/ml ampicillin at 37°C was prepared and added to ^15^N M9 minimal medium the next day. At A600 of 0.95 the expression was induced by the addition of 0.25mM IPTG and incubated at 20°C for 24h (A600 of 3.1). Cell pellets obtained by centrifugation at 5000rpm were stored at −20°C. Cells were solubilized in lysis buffer (20mM TRIS (pH 7.5), 500mM NaCl, 1mM TCEP, 5% glycerol) and disrupted by sonication (Sonopuls from Bandelin) on ice. The sonicated lysate was clarified by centrifugation at 15000rpm for 40min. The supernatant was loaded onto a glutathione-Sepharose-4B affinity column (GE Healthcare) equilibrated with lysis buffer and washed until a stable baseline was obtained. The beads were mixed with TEV protease and incubated at 4°C overnight and washed with lysis buffer. The flow-through was concentrated by centrifugation using an Amicon Ultra 10000 MWCO. The concentrated solution was loaded onto a HiLoad 16/600 Superdex 75pg gel-filtration column (GE Healthcare), equilibrated with 20mM TRIS (pH 7.5), 200mM NaCl, 1mM TCEP. The first major peak (containing TEV) was discarded and the second peak, corresponding to 11kDa in SDS-PAGE analysis, was concentrated to 11.4mg/ml.

### Measurement of the interaction between WRN HRDC and WRN Helicase

^1^H-^15^N SoFast HMQC experiments(52,53) were recorded in 3mm NMR tubes (200 μL filling) at a protein concentration of ^15^N-labeled WRN HRDC of 10 μM ± unlabelled WRN Helicase (residues 517-1093) 20 μM in sample buffer (Tris 20 mM pH7.5, NaCl 150 mM, ATP 250 μM, MgSO4 1.5 mM, D2O 10%). Spectra were recorded on a Bruker Avance III 700 MHz spectrometer equipped with a cryogenically cooled 5mm TCI probe at a 298 K and 256 scans, 128 f1 increments and 2k data points in f2. Total acquisition time was 2 hours.

### HDX MS measurements of WRN in solution

Deuterium labelling: Deuterium labelling was initiated by diluting 3 μL of WRN (17.66μM; 25 mM HEPES and 250 mM NaCl, at pH 7.5) 16-fold in deuterated buffer at room temperature. The labelling reactions were quenched by decreasing the temperature to 0°C and the pH to 2.5 by adding 48 μL of quench buffer. Quench buffer 1 (100 mM potassium phosphate, pH 2.1) was used for the AMP-PNP experiments and quench buffer 2 (4M guanidine hydrochloride, 200 mM potassium phosphate, 200 mM sodium chloride, 50 mM tris (2-carboxyethl) phosphine hydrochloride (TCEP-HCl), pH 2.1) was used for binding experiments where ssDNA was present. Samples were taken at five time points (10 seconds, 1 minute, 10 minutes, 1 hour, 4 hours). The sequence of the ssDNA used in the experiments was as follows: 5’-CCA GGT CGA TAG GTT CGA ATT GGT T - 3’. Complexes with AMP-PNP, ssDNA and ssDNA+AMP-PNP were analysed in a similar way. WRN was incubated with AMP-PNP or ssDNA +AMP-PNP individually. Mixing ratios protein : AMP-PNP of 1 : 100 and protein : ssDNA 1 : 2.5 were used. The protein was allowed to equilibrate with the ligands for 10 min at room temperature before D2O labelling which was allowed to proceed from 10 seconds up to 4 hours for each condition.

LC/MS: Upon quenching, the samples were injected immediately into a Waters nanoACQUITY UPLC equipped with HDX technology. The samples were digested online using a Waters Enzymate™ BEH Pepsin Column (2.1 x 30 mm, 5 μm) at 15 °C. The cooling chamber of the UPLC system which housed all the chromatographic elements was held at 0.0 ± 0.1 °C for the entire time of the measurements. Peptides were trapped and desalted on a VanGuard Pre-Column trap (2.1 mm × 5 mm, ACQUITY UPLC BEH C18, 1.7 μm (Waters, 186003975) for 3 minutes at 100 μL/min. Peptides were then eluted from the trap using a 8%–35% gradient of acetonitrile (with 0.1 % formic acid) over 8 minutes at a flow rate of 40 μL/min, and separated using an ACQUITY UPLC C18 BEH 1.7 μm, 1.0 mm × 50 mm column (Waters, 186002350). The back pressure averaged ^~^7,500 psi at 0 °C and 5% acetonitrile 95% water. The error of determining the deuterium levels was ± 0.15 Da in this experimental setup. To eliminate peptide carryover, a wash solution of (1.5 M guanidinium chloride, 0.8% formic acid and 4% acetonitrile) was injected over the pepsin column during each analytical run. Mass spectra were acquired using a Waters Synapt G2-Si HDMS^E^ mass spectrometer. The mass spectrometer was calibrated with direct infusion of a solution of glu-fibrinopeptide (Sigma, F3261) at 200 femtomole/μL at a flow rate of 5 μL/min prior to data collection. A conventional electrospray source was used and the instrument was scanned at 0.4 scans/second over the range 50 to 2000 m/z with ion mobility engaged. The instrument configuration was the following: capillary voltage 3.2 kV, trap collision energy 4 V, sampling cone 40 V, source temperature 80 °C and desolvation temperature 175 °C. All comparison experiments were done under identical experimental conditions such that deuterium levels were not corrected for back-exchange and are therefore reported as relative.

Peptides were identified using PLGS 3.0.1 (Waters, RRID: SCR_016664, 720001408EN) using three replicates of undeuterated control samples. Raw MS data were imported into DynamX 3.0 (Waters, 720005145EN) and filtered as follows: minimum consecutive products: 2; minimum number of products per amino acid: 0.2. Peptides meeting these filtering criteria were further processed automatically by DynamX followed by manual inspection of all processed data. The relative amount of deuterium in each peptide was determined by subtracting the centroid mass of the undeuterated form of the peptide from the deuterated form at each time point and for each condition. These deuterium uptake values were used to generate uptake graphs and difference maps.

### Data Availability

The crystal structure of WRN was deposited in the Protein Databank PDB ID 6YHR. All HDX MS data have been deposited to the ProteomeXchange Consortium via the PRIDE (54) partner repository with the dataset identifier PXD018910.

## Acknowledgements

The SGC is a registered charity (number 1097737) that receives funds from AbbVie, Bayer Pharma AG, Boehringer Ingelheim, Canada Foundation for Innovation, Eshelman Institute for Innovation, Genome Canada, Innovative Medicines Initiative (EU/EFPIA) [ULTRA-DD grant no. 115766], Janssen, Merck KGaA Darmstadt Germany, MSD, Novartis Pharma AG, Ontario Ministry of Economic Development and Innovation, Pfizer, São Paulo Research Foundation-FAPESP, Takeda, and Wellcome [106169/ZZ14/Z]. MCR is a member of the Boehringer Ingelheim Discovery Research global post-doc program.

The authors would like to thank Diamond Light Source for beamtime (proposal MX19301), and the staff of beamline I03 for assistance with crystal testing and data collection.

## Author contributions

Joseph A. Newman, Angeline E. Gavard, Simone Lieb, Madhwesh C. Ravichandran, Katja Hauer, Patrick Werni, Leonhard Geist, Jark Böttcher, John. R. Engen, Klaus Rumpel, Matthias Samwer, Mark Petronczki and Opher Gileadi, JAN designed, performed and analysed experimental data. AEG, SL, MCR, JRE, KH, KR designed and performed experiments and analysed data. PW and JB prepared WRN HRDC and WRN helicase proteins for HDX measurements and NMR interaction studies performed by LG. JAN, MP and OG supervised the project. JAN, MCR, MP and xxx wrote the manuscript.

