## Supplementary Figs S1-S4 and table S1 for "Crystal Structure of the Werner’s Syndrome Helicase" (from another version of this preprint: version 1, 2020-05-05)

**Supplementary information: Crystal Structure of the Werner's Syndrome Helicase**

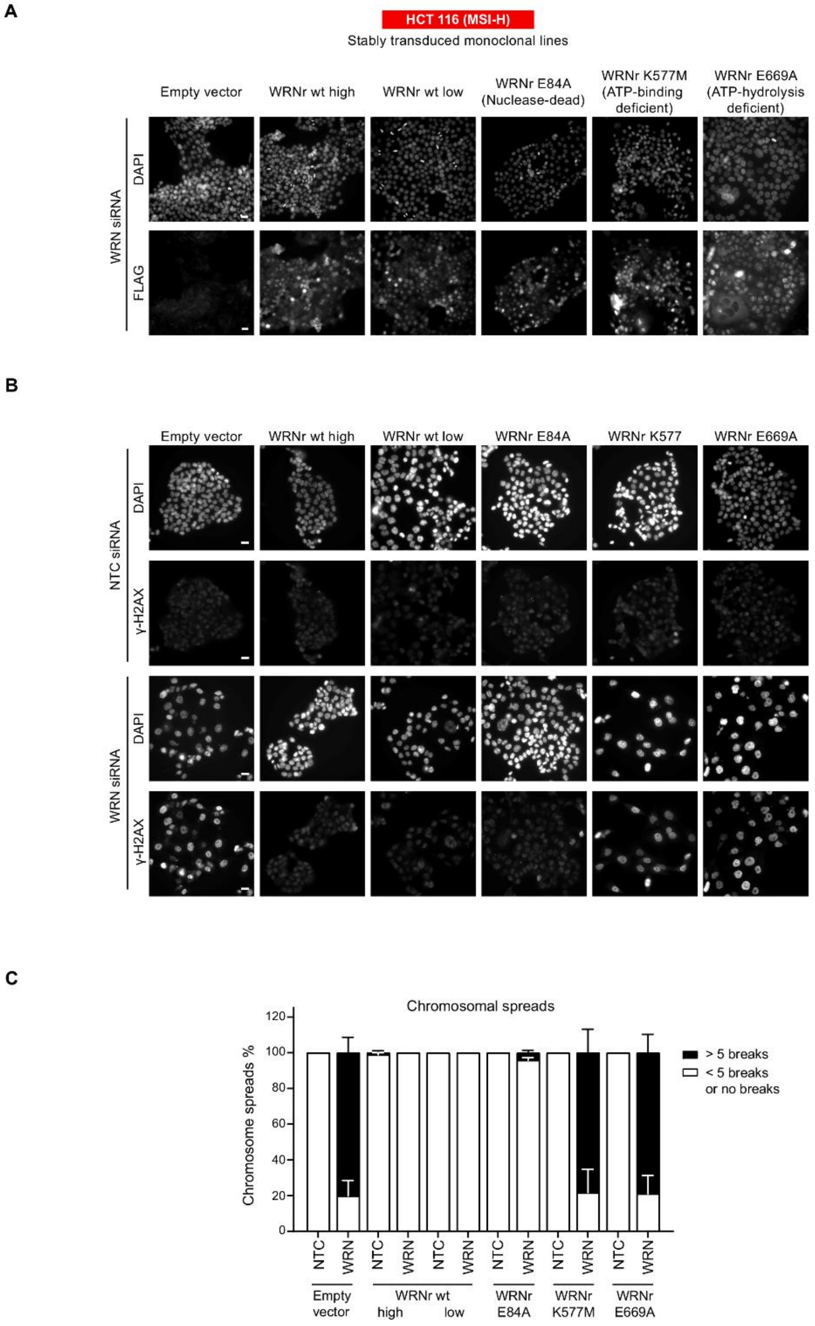


**Figure S1** (A) To monitor cellular expression and nuclear accumulation of the WRNr transgenes an anti-FLAG immunofluorescence analysis was performed in monoclonal cell lines expressing wild-type and mutant variants of WRNr. Scale bar, 20 µM. (B) Representative images of the immunofluorescence analysis of γ-H2AX that was performed 72 hr after siRNA transfection. Scale bar, 20 µM. (C) Mitotic chromosome spread analysis was performed on each of the transgenic cell lines cell lines 72 hr post siRNA transfection that included a 6 hr Nocodazole (1.5 μM) treatment. Quantification was performed under a microscope where each mitotic spread was categorized into less than five (< 5) breaks or more than five (> 5) breaks (n ≥ 28 mitotic spreads). Data values and error bars presented here are the mean and the standard deviation respectively from biological repeats (n = 2).

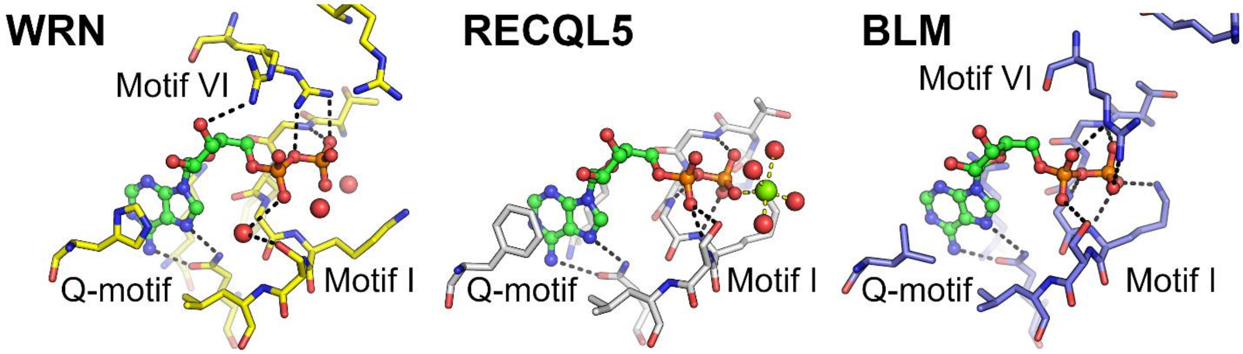


**Figure S2** Comparison of the nucleotide binding modes of WRN (left), RECQL5 (centre) and BLM (right) helicases. The nucleotide in the WRN structure is bound with extensive contacts from motif VI, whilst contacts from motif I are less extensive than in other RecQ helicase structures.

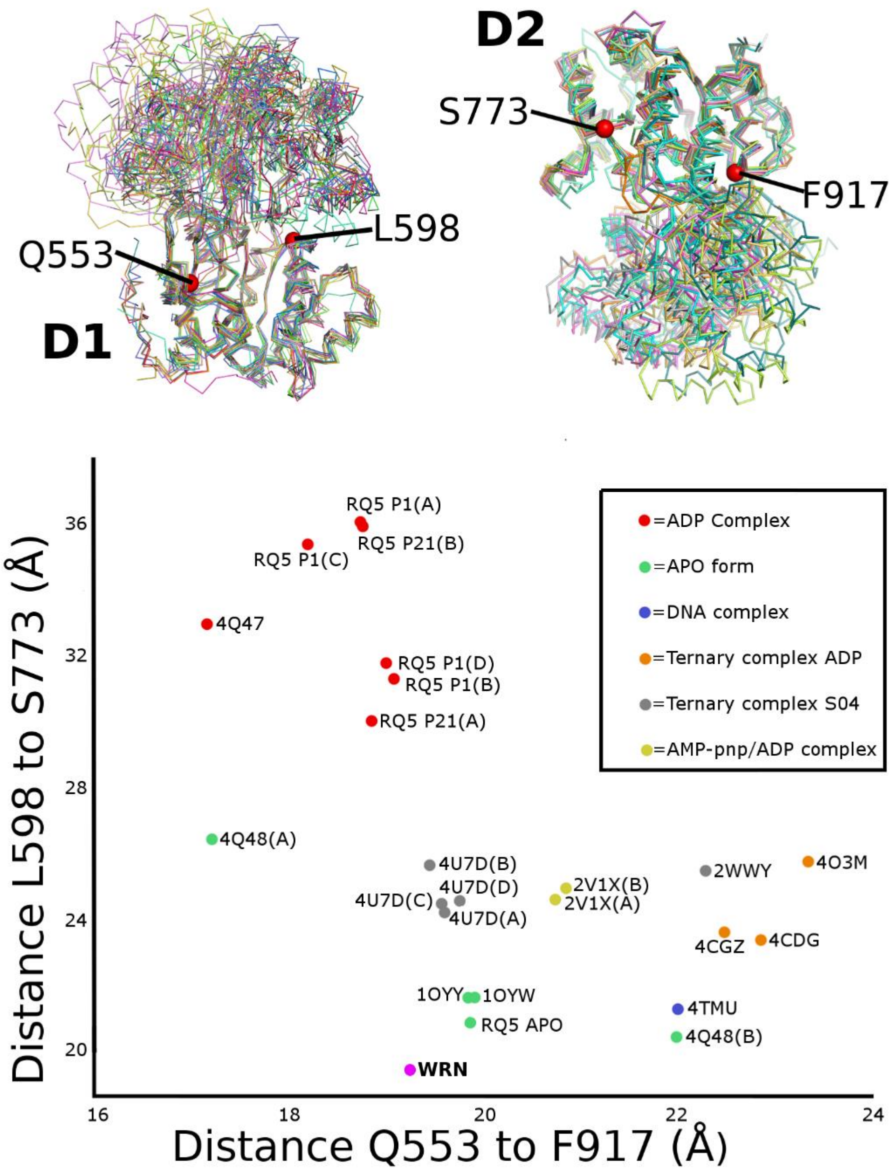


**Figure S3** Global comparison of RecQ family relative domain positioning of D1 and D2. The upper panel shows a structural superposition of selected RecQ family structures on the basis of the D1 domain (left) or D2 domain (right). Invariant points used to define the positioning of the two domains are depicted as red spheres with numbering according to the WRN sequence. The lower panel shows a 2D vector diagram of distances between the vector pairs. The WRN structure occupies a distinct position at the bottom of the plot with a conformation even more compact that the APO form structures.

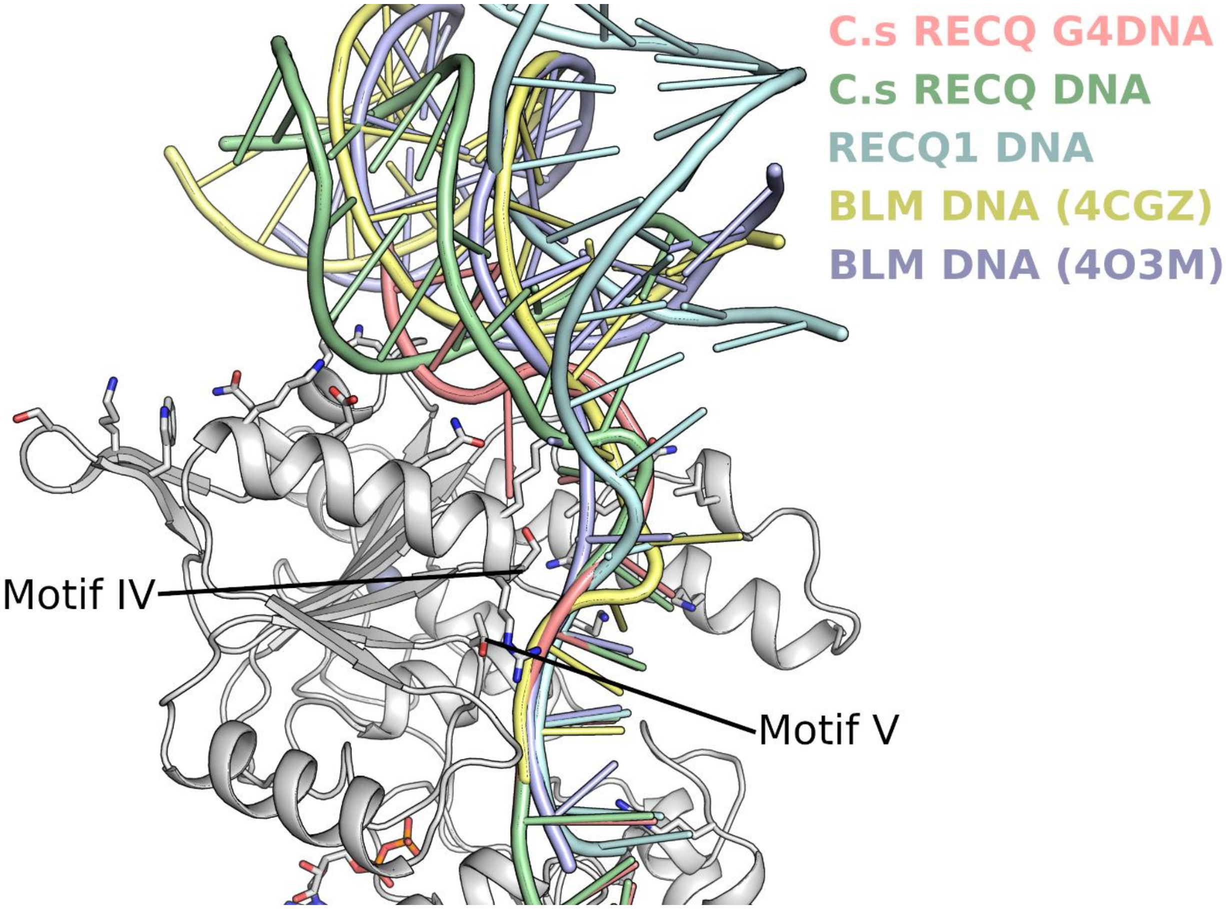


**Figure S4** Comparison of current RecQ family DNA complex structures. The double stranded part of the DNA is positioned variably with respect to the D2 domain whilst the single stranded 3’ overhang can be seen to converge an makes conserved interactions with helicase motifs IV and V.

**TABLE S1. HDX Data Summary and list of experimental parameters**

| **Data Set** | WRN +AMP-PNP | WRN +ssDNA +AMP-PNP |
|---|---|---|
| **States analyzed** | State 1: WRN alone; State 2: WRN + AMP-PNP | State 3: WRN alone; State 4: WRN +ssDNA +AMP-PNP |
| **HDX reaction details** [a-c] | Final $D_2O$ concentration = 93.8%, $pH_{read}$ = 7.1[a] | |
| | Quench buffer 1[b] | Quench buffer 2[c] |
| **HDX time course** | 0.167, 1, 10, 60, 240 minutes | 0.167, 1, 10, 60, 240 minutes |
| **HDX controls** | 4 undeuterated controls:<br>3 for state 1, 1 for state 2 | 2 undeuterated controls:<br>1 for state 3, 1 for state 4 |
| **Back-exchange** | 25-30% | |
| **Number of peptides** | 94 followed; 155 identified | 62 followed; 127 identified |
| **Filtering parameters** | 0.3 products per a.a., 2 consec~~t~~utive products,<br>10ppm error | 0.3 products per a.a., 2 consec~~t~~utive products,<br>10ppm error |
| **Sequence coverage** | 98.8% | 73.2% |
| **Average peptide length** | 13.3 | 10.7 |
| **Redundancy** | 3.00 | 2.17 |
| **Replicates** | 1 technical for each state | 1 technical for each state |
| **Repeatability** | +/- 0.15 relative Da | |
| **Significant differences** | > 0.5 Da | |

[a] 16-fold dilution with labeling buffer [25 mM HEPES pD 7.5, 150 mM NaCl, 99.9% $D_2O$]. 1:1 dilution with quench buffer 1 or 2

[b] Quench buffer 1: 100 mM potassium phosphate, $H_2O$, pH 2.1

[c] Quench buffer 2: 4 M guanidine hydrochloride, 200 mM potassium phosphate, 200mM sodium chloride, 50mM tris(2-carboxyethyl) phosphine hydrochloride (TCEP-HCl), $H_2O$, pH 2.1
